# The nucleus spatially partitions mechanoprotection and mechanosensing

**DOI:** 10.1101/2025.08.25.672162

**Authors:** Ronan Bouzignac, Amandine Palandri, Lauriane Fritsch, Tatiana Merle, Amal Zine el Aabidine, Thomas Mangeat, Mélanie Gracia, Cyril Andrieu, Pablo Garcia-Idieder, Antonio Trullo, Martine Cazales, Simon Marques-Prieto, Virginia Pimmett, Mélissa Costes, Christian Rouviere, Mounia Lagha, Magali Suzanne

## Abstract

Mechanical forces shape cell behavior, but how the nucleus integrates its roles as genome protector and mechanosensor remains unresolved. Here, we dissect the spatiotemporal control of transcriptional response to forces during epithelial-to-mesenchymal transition (EMT). By combining genome-wide approaches, *in vivo* nuclear micromanipulation, and quantitative live imaging, we uncover two key properties of nuclear mechanosensing. First, mechanically induced changes in chromatin accessibility and transcription are spatially patterned within the nucleus: genes closest to deformations appear protected, revealing a local shielding mechanism associated with chromatin closure. Second, relocating a gene between chromatin environments is sufficient to alter its mechanosensitivity, challenging the concept of inherently force-sensitive genes. These findings demonstrate unexpected spatial partition of nuclear response to forces.

## Introduction

The epithelio-mesenchymal transition (EMT) is a cellular process crucial to form different cell layers during development. Importantly, EMT is also associated with increased drug resistance and invasiveness in cancer ^1^. The transition from epithelial to mesenchymal status notably includes numerous intermediate steps of force generation ^2^, which coincide with a critical transcriptional shift. Importantly, forces can be sensed by the nucleus. During cell differentiation, it was shown that the nucleus can respond to forces in two opposite manners, acting as a shield to protect nuclear content from external mechanical forces ^3,4^ or as a ‘transducer’ to convert mechanical stimuli into nuclear biochemical signals, a process known as mechanotransduction ^5,6^. However**, how the balance between nuclear mechanoprotection and mechanotransduction is established remains largely unexplored in the complex environment of a living organism and totally unknown during EMT.**

The temporal evolution of nuclear responses to mechanical challenges is also an open question. Recent work proposed that different responses to mechanical stresses may reflect distinct temporal phases of cellular adaptation ^7^, highlighting the importance of considering temporal resolution in a mechanotransduction response. Mechanical forces are transmitted to the nucleus within seconds or milliseconds ^8–10^. This fast transmission constitutes a hallmark of mechanotransduction, operating far more rapidly than any biochemical signaling pathway ^8,11^. However, transcriptional responses to mechanical cues have thus far only been apprehended at the scale of minutes or hours to days ^7,12–15^. **Thus, how fast transcriptional response to mechanical challenges is taking place remains an open question.**

To address how EMT nuclei respond to mechanical cues *in vivo*, we used *Drosophila* mesoderm invagination, a well-established model of EMT, and combined quantitative imaging approaches, genetics, nuclei micromanipulation with optical tweezers, compression experiments and genome-wide assays. We revealed a rapid, spatially patterned mechanical response during EMT, orchestrated along the apico-basal nuclear axis, with a protected apical region, and a more mechanically sensitive medio-basal region.

## Results

### Nuclei undergo stereotyped deformations at the onset of EMT

During the various steps constituting an EMT, cells generate different types of forces ^2^. To assess how the nucleus responds to these mechanical stresses, we analyzed nuclear shape using Lamin Dm0 staining during *Drosophila* mesoderm invagination. Prior to mesoderm invagination and associated EMT, nuclei exhibit a regular and elongated shape along the apico-basal axis (Fig. 1A). In contrast, during invagination, stereotypical apical deformations were observed (Fig. 1A-B). To account for inter-nuclear variability, we analyze nuclear shape at a single nucleus level, using the distance from the apical surface of the embryo (depth) as a proxy of EMT progression (Fig. 1B). This analysis revealed a temporal correlation between apical nuclear deformation and basal repositioning (Fig. 1C), indicating that nuclei are progressively deformed during apical constriction.

**Figure 1:**
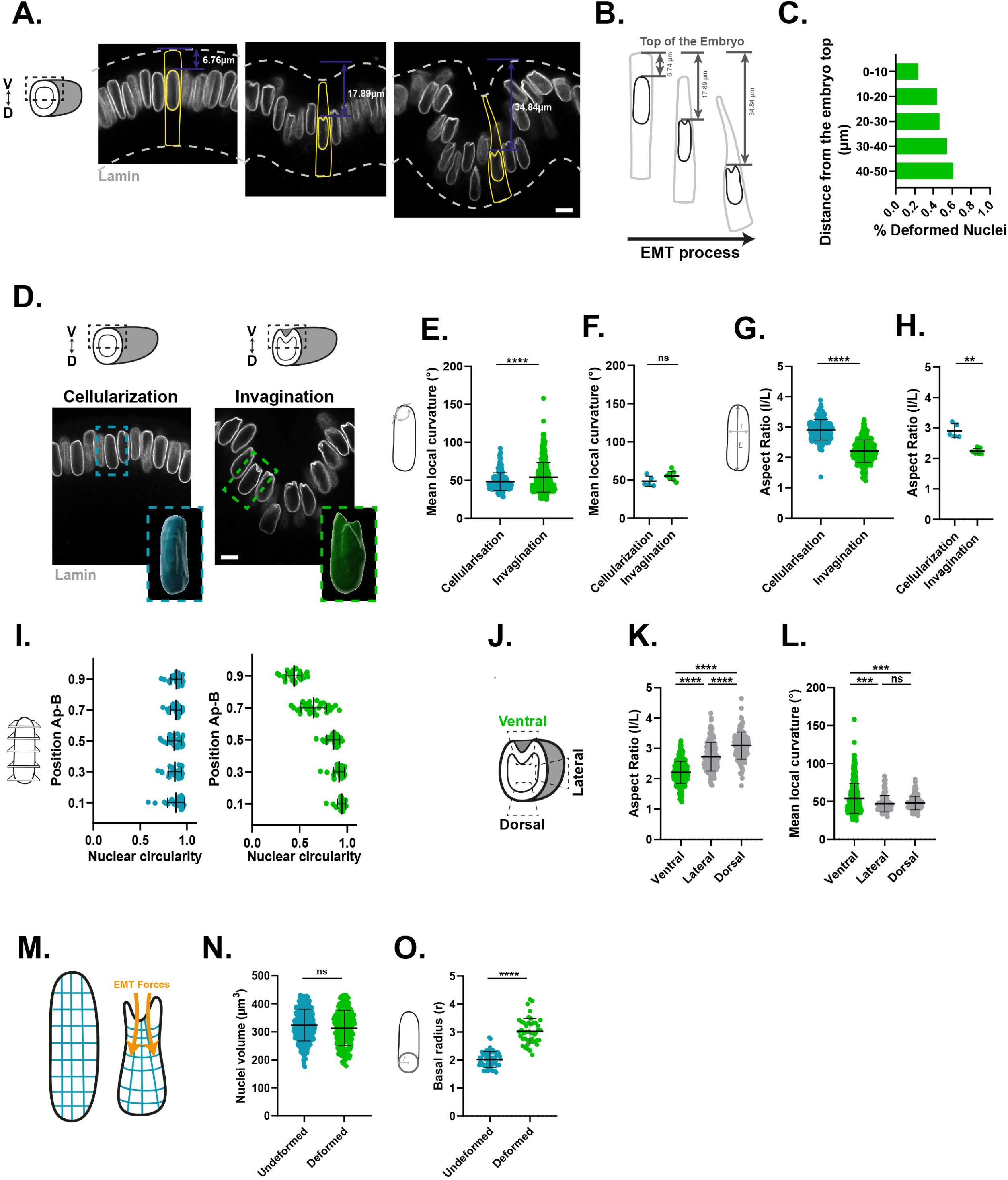
Nuclei undergo stereotyped apical deformations during EMT. Related to Fig. S1. **A.** Transverse sections of *wt* embryos stained for Lamin Dm0 during invagination. Nuclear and cellular outlines are indicated in yellow; dashed lines mark the apical and basal tissue surfaces. Nuclear depth was measured from the apical surface. Note the heterogeneity in nuclear depth across the embryo. **B.** Schematic of EMT progression showing the relationship between apical constriction, nuclear repositioning and nuclear deformation. **C.** Quantification of the percentage of deformed nuclei in *wt* embryos as a function of their depth within the epithelium (N=13 embryos, n= 629 nuclei). **D.** Transverse sections of embryos stained for Lamin Dm0 during cellularization (pre-EMT, left) and invagination (EMT, right). Insets show selected nuclei in 3D. **E-F.** Quantification of the nuclear local curvature during cellularization (pre-EMT) and invagination (EMT) (cellularization: N=5, n=198; invagination: N=8, n=431), per nucleus (E), or per embryo (F). **G-H.** Quantification of the nuclear aspect ratio during cellularization (pre-EMT) and invagination (EMT) (cellularization: N=5, n=198; invagination: N=8, n=431), per nucleus (G), or per embryo (H). **I**. Quantification of nuclear circularity at different positions along the apico-basal axis of the nuclear envelope during cellularization (pre-EMT; left) and invagination (EMT; right) (N=3, n=30). **J.** Schematic of an embryonic transverse section showing ventral, lateral and dorsal regions used for the analysis in K and L. **K-L.** Measurement of nuclear aspect ratio (K) and nuclear local curvature (L) during invagination across the indicated regions (ventral: N=8, n=431; lateral: N=3, n=159; dorsal: N=3, n=149). **M.** Schematic representation of a nuclear cross-section illustrating how EMT-generated forces deform both the apical and basal regions of the nucleus. **N.** Quantification of nuclear volume in undeformed and deformed nuclei (undeformed: N=13, n=406; deformed: N=13, n=223). **O.** Quantification of nuclear basal radius in undeformed and deformed nuclei of *wt* embryos (undeformed: N=4, n=60; deformed: N=4, n=45). Statistical significance was assessed by the Mann−Whitney test (Fig. 1 F,G, H, K, N, O), Kolmogorov-Smirnov test (Fig. 1E(left), L) Statistical significance was assessed by the Mann−Whitney test (Fig. 1E, F, I, L, M), Kolmogorov-Smirnov test (Fig. 1E(left), J), (**\***=p<0.05; **\*\***=p<0.01; **\*\*\***=p<0.001; ****p<0.0001). N=embryos, n=nuclei. Scale bar = 5 µm.

When comparing nuclei pre-EMT (cellularization) or during EMT (invagination) (Fig. 1D), clear differences in local curvature of the nuclear envelope (Fig. 1E-F), aspect ratios (Fig. G-H) and circularity emerge (Fig. 1I). Interestingly, these deformations are restricted to the apical part of the nuclei (Fig. 1A, D, I). Moreover, they are more prevalent in the mesoderm compared to lateral and dorsal regions (Fig. 1J-L). In addition, upon deformation, the nuclear volume stays constant, but its content shifts basally, increasing the basal diameter (Fig. 1M-O). Thus, collectively, these results show that apical deformation is only the tip of the iceberg, while EMT forces may exert a broader mechanical influence on the entire nucleus at the onset of EMT.

Interestingly, we found that these deformations coincide with changes in the physical properties of the nucleus. By trapping lipid droplets (naturally present in the embryo and used as internal probes), we locally indented a target nucleus (see Fig. S1A, and Fig. S1B for a schematic representation). We found that wt nuclei were significantly softer at the onset of EMT (beginning of invagination) compared to 10 minutes earlier (during late cellularization) (Fig. S1C) and relaxation time slightly increased (Fig. S1D), revealing temporal changes in nuclear viscoelasticity.

In *Drosophila* embryos, prior works using static approaches suggested that *twi* and *sna* expression were sensitive to mechanical cues ^19,20^. We therefore sought to test whether the nascent transcription of *sna* and *twi* could respond to these early EMT forces. We used smiFISH labeling and scored two metrics: the number of active transcriptional site (TS) per nucleus or transcriptional activation and the sum of TS intensity per nucleus or transcriptional level. We found that sna transcription was mainly unchanged between deformed and undeformed nuclei (Fig. S1E-G), while *twi* transcription was significantly increased in deformed wt nuclei, both in terms of transcriptional activation and transcriptional level per nucleus (Fig. S1E’-G’).

Thus, nuclear deformations occurring at the onset of EMT coincide with nuclear envelope softening and changes in *twi* expression level.

### Controlled Micromanipulation of the Nucleus in living embryos Reveals Direct Mechanical Impact on Transcription

We next asked whether a local mechanical stress applied directly on the nuclear envelope would be sufficient to impact gene transcription during EMT. To test this, we used optical tweezer to apply a local indentation at precise location of the nuclear envelope (Fig. 2A-B). To monitor the transcriptional response in real time, this approach was coupled to the MS2/MCP labeling system, which enables live imaging of transcription by recruiting fluorescently tagged MS2 coat protein (MCP) to MS2 RNA stem loops inserted into the nascent transcript (see Methods). We first followed the response of endogenous *sna* (with a *sna*-MS2 crispr allele^22^) to local indentations generated at the level of the *snail* locus. Such indentation failed to induce any change in *sna* transcriptional activity, regardless of the distance between the indentation and the TS (Fig. 2C-D, Movie S1). In contrast, performing the same experiment on another region of the nuclear envelope, at the level of the endogenous *twi (twi-MS2* Crispr KI^23^) resulted in a rapid and significant increase in transcriptional signal observed within 8-10 seconds after applying the local force (Fig. 2C’-D’). This effect demonstrates that a direct mechanical perturbation of the nuclear envelope is sufficient to stimulate *twi* transcription in *Drosophila* embryos. Of note, because the deformations were induced very locally, the response was strictly local: only TS located within ∼2.5 µm of the indentation showed increased transcription, while more distant TS remained unaffected (Fig. 2C’-D’, Movie S2). By applying controlled forces directly to the nuclear envelope in living embryos, these experiments demonstrate the immediate impact of mechanical stress on the transcriptional response. They also reveal a differential response of *sna* and *twi* genes to mechanical stresses.

**Figure 2:**
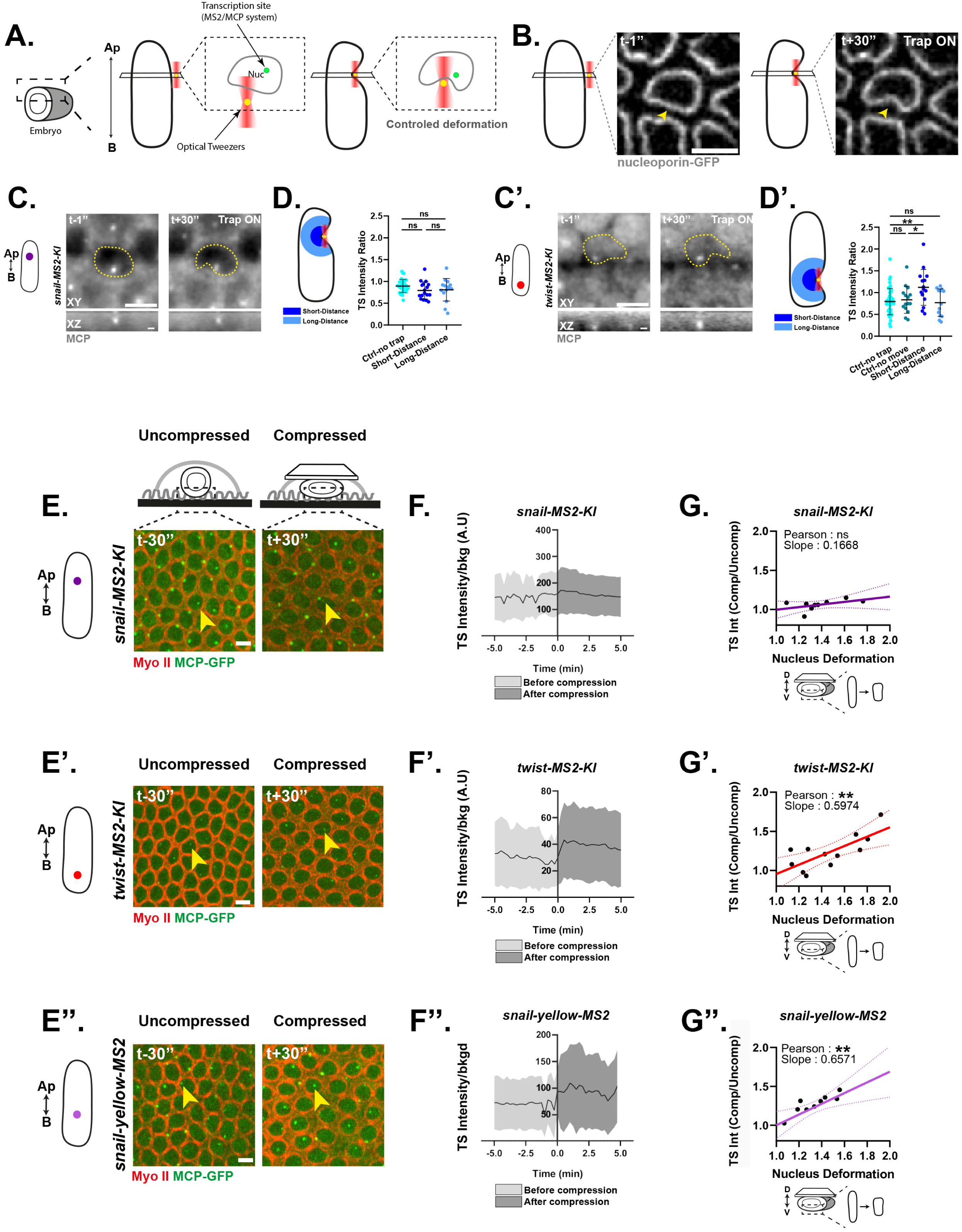
Gradual and immediate response of *twist* but not *snail* to mechanical challenges. Related to Fig. S2 and Movies S1-5. **A.** Schematic representation of the optical tweezer/MS2 experiment. **B.** Representative time-lapse images of the nuclear envelope (Nup107–GFP) before (left) and during (right) local nuclear deformation. Yellow arrow indicates the initial trap position. **C-C’.** Representative images from live MS2/MCP experiments showing TS before (left) and during (right) an induced localized nuclear deformation (top: XY axis; bottom: XZ axis, scale bar = 1µm). C: *snail-MS2-KI*; C’: *twist-MS2-KI*. Yellow dashed lines indicate the estimated nuclear boundaries, based on brightfield images (data not shown). **D-D’.** (left) Schematic of the quantification method. TS within 2.5 µm of the initial trap position were classified as “short distance”, and those farther away as “long distance”. (right) Change in TS intensity before versus after local deformation. “Ctrl-no trap” denotes adjacent unmanipulated nuclei, and “Ctrl-no move” nuclei in which lipid droplets were trapped nearby but not displaced against the nuclear envelope. (*snail*: N=7, Ctrl-no trap =33, Short distance=17, Long distance=15; *twist*: N=17, Ctrl-no trap =43, Ctrl-no move = 15, Short distance=15, Long distance=13). **E-E’-E’’.** Time-lapse snapshots from the compression experiment shown before (left) and after (right) compression for *snail-MS2-KI* (E), *twist-MS2-KI* (E’) and *snail-yellow-MS2* BAC (E’’). Yellow arrows indicate the same transcription site before and after compression. MyoII is shown in red, MCP-eGFP (TS) in green. **F-F’-F’’.** Representative curves of mean normalized TS intensities over time during compression of *snail-MS2-KI* (F), *twist-MS2-KI* (F’), *snail-yellow-MS2* BAC (F’’). **G-G’-G’’.** Curves showing the variation in TS intensity (ratio before *vs* after compression) in relation to the level of nuclear deformation (nuclear aspect ratio variation, see Fig. S2) in *snail-MS2-KI* (G), *twist-MS2-KI* (G’) and *snail-yellow-MS2* BAC (G’’) embryos (N=9, 12, 9 respectively). Statistical analyses: Mann–Whitney test (Fig. 2D, D’) and Pearson correlation (Fig. 2F-G, F’-G’, F’’-G’’) (p<0.05 =*; p<0.01=**; p<0.001=***; ****p<0.0001). N=embryos, n=nuclei. Scale bar = 5 µm.

### Gradual and immediate response of *twist* but not *snail* to mechanical challenges

To complement the optical-tweezer experiments, we developed a simpler orthogonal approach: gently compressing living embryos by placing a coverslip on top of them. Compression was applied to living embryos at pre-EMT stage while monitoring transcriptional activity in real time using the MS2/MCP system (as exemplified in Fig. S2A-D). For each experiment, the nuclear aspect ratio was used as a readout for the level of compression applied (Fig. S2E-G). Compression induces both embryo flattening and nuclear compression (Fig. S2H-J). Importantly, the induced nuclear deformation was slightly higher but of the same order of magnitude as that observed during natural mesoderm invagination (Fig. S2F). Upon compression, *sna* nascent transcription remained stable (Fig. 2E-F, Movie S3), independently of the level of compression applied (Fig. 2G). This confirms that *sna* transcription is not directly responsive to mechanical forces, as observed previously with optical tweezers, at least in the range of forces applied in these experiments. In contrast, compression triggered a rapid and substantial increase (∼ 40%) in *twi* transcriptional activity (Fig. 2E’-F’, Movie S4). The intensity of transcriptional activation correlated linearly with the extent of nuclear deformation, indicating that greater nuclear deformation elicits a stronger *twi* transcriptional response (Fig. 2G’). Consistent with the optical tweezer experiments (Fig. 2C’-D’), these results also revealed a very rapid transcriptional response to force application (Fig. 2F’).

Given the absence of response at the endogenous *sna* locus, we examined a second *sna*transcriptional reporter, a *sna-yellow-MS2* BAC reporter inserted in another chromosome, which includes the whole regulatory regions of *sna* ^24^. Surprisingly, this BAC reporter exhibited a rapid yet gradual transcriptional response to global compressive forces, similar to that observed for *twi* (compare Fig. 2E’’-G’’ to Fig. 2E’-G’, and see Movie S5). Since this construct is localized at a different locus than the endogenous *sna*, this result suggests that the transcriptional response to forces could be shaped by the local chromatin environment rather than being an intrinsic property of the gene.

### The nuclear microenvironment shapes the response to mechanical forces

Given the distinct responses of endogenous *sna*, *twi*, and the *sna*-BAC reporter to compression, together with their localization in different nuclear regions, we hypothesized that nuclear positioning and the associated chromatin environment determine their differential mechanosensitivity.

To directly test the effect of nuclear positioning, we examined the response to global compression of the same MS2 reporter inserted at different apico-basal nuclear positions (Fig. 3A-B). In order to survey various apico-basal positions, we took advantage of the available PhiC31 landing sites ^25^ and the stereotypical Rabl chromosomal configuration of the *Drosophila* nuclei at this stage of development ^26^ (Fig. 3B). The Rabl organization, consists in the clustering of centromeres and telomeres at opposite nuclear poles ^27^. We consistently observed a strong correlation between the position of genes along the apico-basal axis of the nucleus (using the TS from smiFISH experiments as a proxy for gene position) and their distance from the centromere for genes on chromosomes II and III (Fig. S3A-B). This correlation curve allowed us to infer apico-basal positions for any given genomic coordinate (see Methods). The same *sna*-MS2 (this study) or *twist*-MS2 ^28^ reporter transgene (including minimal regulatory regions, see Methods) was inserted in distinct location along the apico-basal axis of the nucleus. Interestingly, for both *sna* and *twi* transgenic reporters, we found that the response to compression was clearly dependent on the subnuclear position of the MS2 reporter, with the apical most positions being the least responsive to forces, while basal positions strongly respond to compression (Fig. 3C-E). For both *sna*- and *twi*-MS2 constructs, the response was gradual, as shown by the linear increase of TS intensity with nuclear deformation (Fig. 3C-E). Together, these experiments highlight nuclear position as an important determinant of transcriptional mechanosensitivity: basal relocation of *sna* enhances its responsiveness to mechanical stimulation, whereas apical relocation of *twi* reduces it. These findings suggest that the nuclear microenvironment can strongly modulate the mechanosensitivity of a given regulatory sequence. However, nuclear positioning alone does not fully determine the transcriptional response to mechanical forces. Indeed, when comparing sna-MS2 and twi-MS2 inserted at the same medial genomic position (vk33), we observed distinct responses to compression: while twi-MS2 remained unresponsive, sna-MS2 exhibited a strong, graded response (Fig. 3C). Thus, our results suggest that transcriptional mechanosensitivity arises from the interplay between regulatory sequence identity and nuclear positioning, with the nuclear microenvironment modulating the intrinsic mechanical responsiveness of a given sequence.

**Figure 3:**
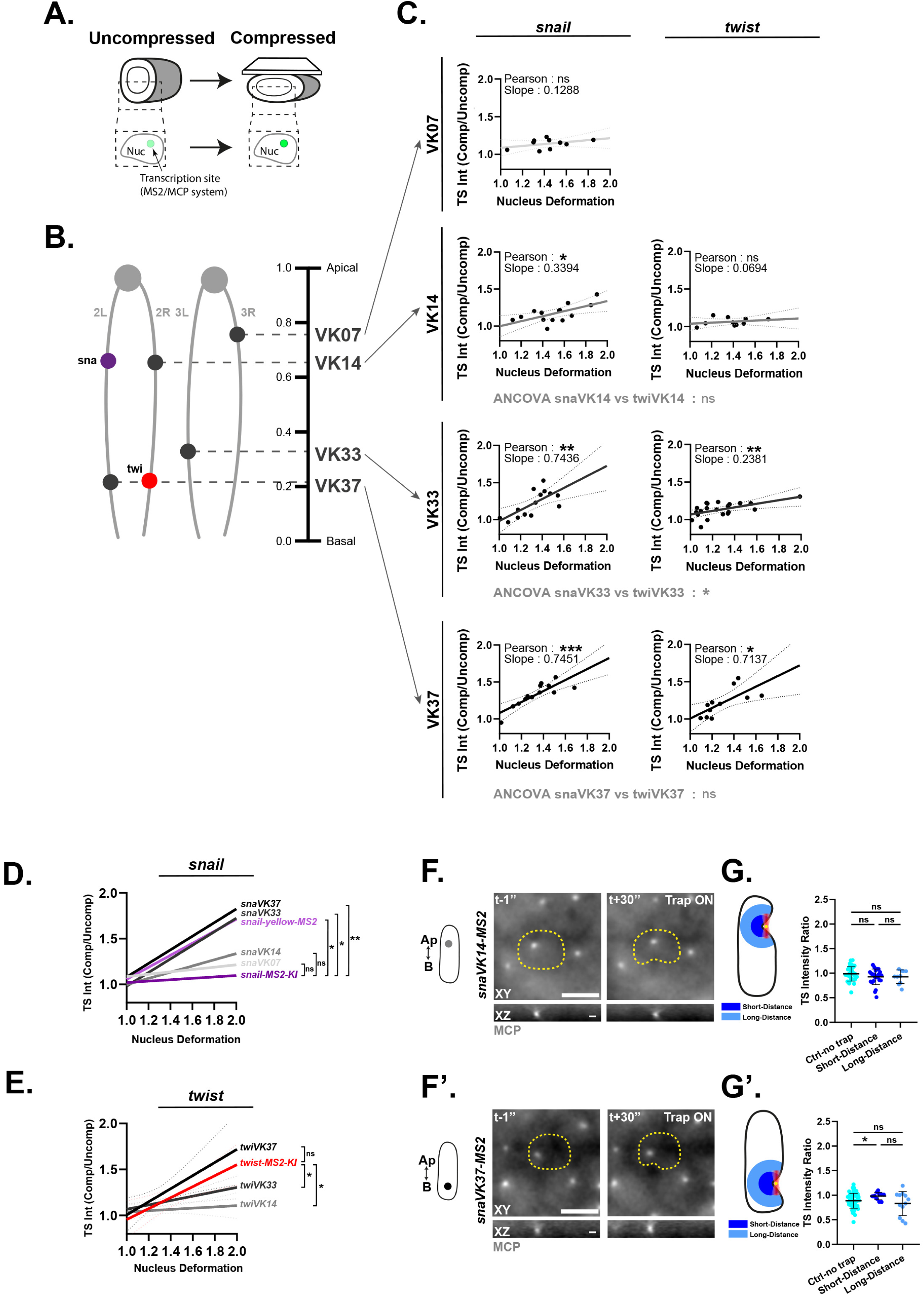
The nuclear microenvironment shapes the response to mechanical forces. Related to Figure S3. **A.** Schematic of transcriptional recording using the MS2/MCP system in compressed living embryos (see Fig. S2 for details). **B.** Schematic showing the chromosomal organization (left) and inferred apico-basal nuclear position (right) of the MS2 reporters analyzed. **C.** Curves showing changes in TS intensity (ratio before vs after compression) as a function of nuclear deformation (change in nuclear aspect ratio; see Fig. S2) for *sna*-MS2 transgenes: *snaVK07, snaVK14, snaVK33* and *snaVK37* (N = 10, 14, 14 and 13, respectively), and *twi*-MS2 transgenes: *twiVK14, twiVK33* and *twiVK37* (N = 10, 21 and 11, respectively). **D, E**. Curves from all compression experiments shown in C and Fig. 2G–G′′, pooled for all *sna*-MS2 (D) or *twi*-MS2 (E) reporters. Statistical differences between curve slopes are indicated. **F-F’.** Representative images from live MS2/MCP experiments showing TS before (left) and during (right) an induced localized nuclear deformation (top: XY axis; bottom: XZ axis, scale bar = 1µm). H: *sna-MS2* at vk14; I: *sna-MS2* at vk33. Yellow dashed lines indicate the estimated nuclear boundaries, based on brightfield images (data not shown). **G-G’.** (left) Schematic of the quantification method. TS within 2.5 µm of the initial trap position were classified as “short distance”, and those farther away as “long distance”. (right) Change in TS intensity before versus after local deformation. “Ctrl-no trap” denotes adjacent unmanipulated nuclei, and “Ctrl-no move” nuclei in which lipid droplets were trapped nearby but not displaced against the nuclear envelope. Statistics: *vk14* N=17, Unmanipulated=35, Short distance=25, Long distance=17; *vk33*: N=11, Unmanipulated=43, Short distance=15, Long distance=13). Statistical analyses: Mann–Whitney test (Fig. 3G, G’), ANCOVA (Fig. 2B, D, E), and Pearson correlation (Fig. 2C), (**\***=p<0.05; **\*\***=p<0.01; **\*\*\***=p<0.001; ****=p<0.0001). N=embryos. Scale bar = 5 µm.

However, this differential response along the apico-basal axis could reflect regional variation in force transmission across the nucleus. To test this, we used optical tweezers to impose controlled deformations at distinct positions along the apico-basal axis of the nuclear envelope. Because Lamin concentration varies along the nuclear envelope (Fig. S3C-D) and heterochromatin density increases upon nuclear deformation (Fig. S3F-H), we first tested for regional differences in nuclear stiffness. We detected no significant variation along the lateral nuclear envelope (Fig. S3E).

Next, we tested the direct transcriptional output to the same local force applied at two distinct nuclear locations, using different insertions of the *sna*-MS2 construct (apico-medial: vk14= 0,65, basal: vk37 = 0,2). When apically positioned, *sna*-MS2 appeared insensitive to the applied force (Fig. 3F-G). In contrast, moving the same reporter to a basal nuclear position was sufficient to render it responsive to an identical mechanical stimulus (Fig. 3F’-G’). Thus, the transcriptional response to force can be tuned by changing the nuclear position of the same regulatory sequence. These findings support a model in which transcriptional mechanosensitivity is determined primarily by the local chromatin environment.

### Mechanically induced changes in chromatin accessibility are spatially patterned

To determine whether the spatially dependent mechanosensitivity observed for *sna* and *twi* extends genome-wide, we employed a genome-wide approach combined with bulk compression to directly manipulate mechanical forces. Cellularized embryos were manually sorted, aligned on a slide, and subjected to uniform compression for 5 min, inducing mild but reproducible nuclear deformation (Fig. S2H-J). ATAC-seq was then performed on compressed and uncompressed wild-type (*wt*), using *D. yakuba* embryos as a spike-in for normalization. This method allowed us to test the impact of forces on the chromatin landscape. We identified numerous regions with altered chromatin accessibility in response to applied forces (applied naturally during invagination or by compression) (See Methods).

The first observation was that compression predominantly induces chromatin closure (Fig. 4A). Using the Rabl configuration to infer the nuclear positions of genomic regions (Fig. S3A-B), we found that this response was concentrated in the most apical region (Fig. 4A). Interestingly, a similar apical chromatin closure was observed at the onset of invagination, suggesting that chromatin closure may represent a physiological response to forces during EMT (Fig. 4B). This regionalization was further confirmed by the clustering of the most significantly affected regions upon compression or during invagination, as shown in Fig. 4C. We observed two clusters, showing a clear spatial segregation along the apico-basal axis of the nucleus. Cluster 2, comprising regions with reduced chromatin accessibility in response to force application (either endogenous forces generated during invagination or externally applied compression) was strongly enriched in the apical nuclear region (Fig. 4D), consistent with previous observations (Fig. 4A-B). By contrast, regions in cluster 1 showed a moderate decrease in accessibility upon compression and increased accessibility during invagination (Fig. 4C). These regions were predominantly localized to the medio-basal region of the nucleus (Fig. 4D). These experiments reveal that the nucleus exhibited a clear spatial regionalization, with an apical region showed a robust reduction in chromatin accessibility upon force application. This apical condensation of chromatin, occurring at the onset of gastrulation, could also be visualized by high-resolution imaging of Histone2B (Fig. 4E-F).

**Figure 4:**
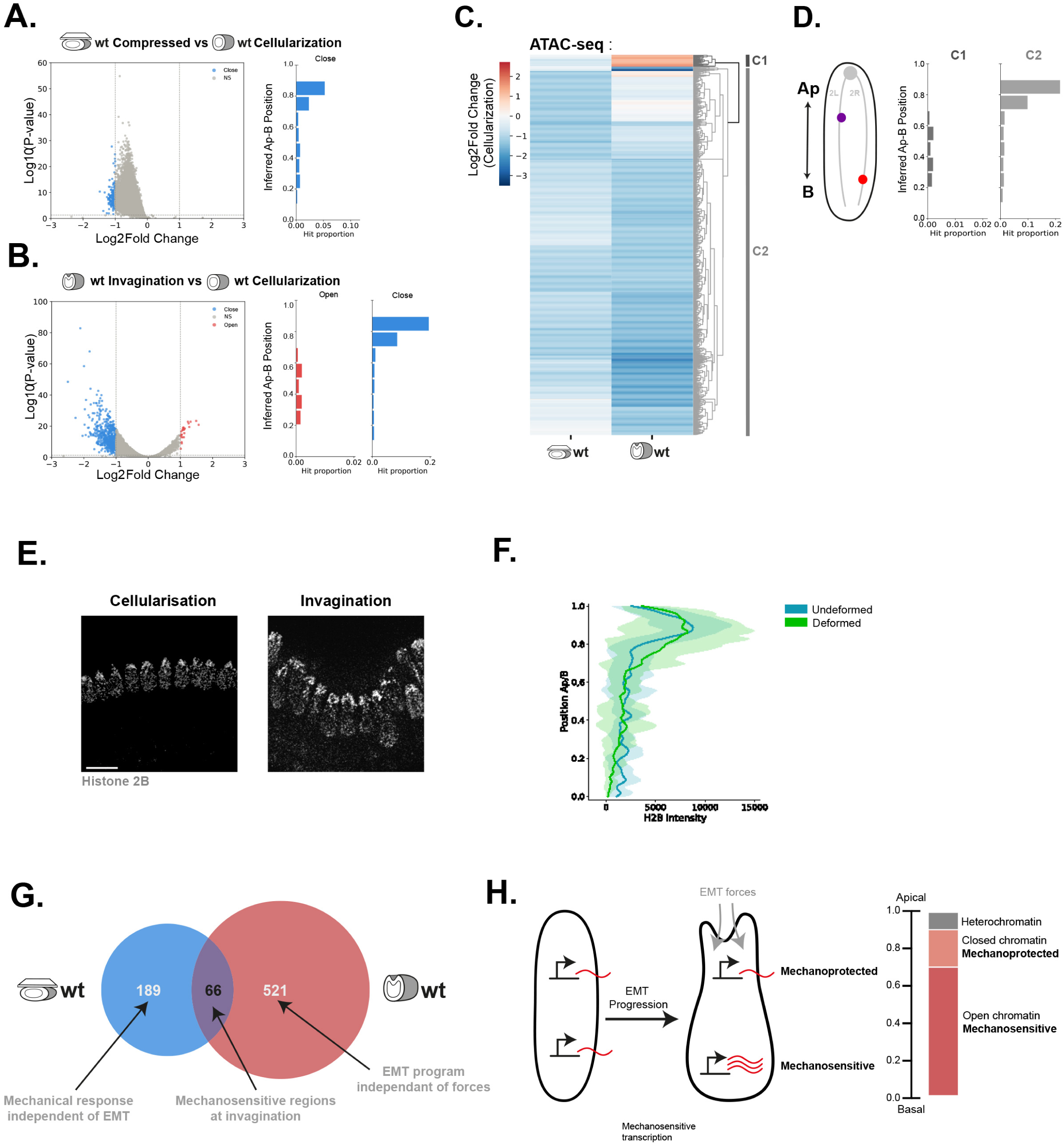
Mechanically-induced changes in chromatin accessibility and transcription are spatially patterned genome-wide. **A-B**. (left) Volcano plot of regions showing differential chromatin accessibility upon compression (A) or during invagination (B) in wt embryos. (right) Distribution of regions with significant changes in chromatin accessibility along the apico-basal nuclear axis. Nuclear positioning was estimated from centromeric distance using the correlation shown in Fig. S3 (see Methods). **C.** Clustering analysis of regions with the most significant differences in chromatin accessibility in *wt* embryos with/without compression and before and during invagination, as measured by ATACseq data. Genes were clustered based on their log2 fold change (compressed vs. uncompressed or invagination vs. cellularization) and grouped by similar chromatin patterns. High ratios (>1) indicate more open chromatin in compressed or invaginating embryos, while low ratios (<1) indicate less accessible chromatin relative to the uncompressed or cellularized condition. **D.** Spatial distribution of regions from the two clusters identified in C along the apico-basal nuclear axis. **E.** Cross-sections of His2Av-mRFP embryos imaged by random illumination microscopy for super-resolution imaging ^39^, showing the distribution of condensed chromatin during cellularization and invagination. **F.** Quantification of His2Av-mRFP signal intensity along the apico-basal axis of the nuclei during cellularization and invagination (cellularization: N=3, n=42; invagination: N=3, n=45). **G.** Venn diagram comparing mechanically induced changes in chromatin accessibility and those occurring during EMT. **H.** Schematic of the polarized transcriptional response to mechanical forces during EMT. Basally positioned genes preferentially respond to nuclear deformation, whereas apical genes are less responsive, potentially owing to increased chromatin compaction.

Importantly, this analysis allowed us to disentangle the effects of mechanical forces on chromatin structure from those associated with EMT commitment. We identified chromatin regions that responded directly to compression, as well as regions that were remodeled during EMT, independently of a mechanical input (Fig. 4G). Notably, only 10% of the regions exhibiting differential chromatin accessibility during EMT were affected by compression. This limited overlap may partly reflect the inability of externally applied compression to fully recapitulate the endogenous mechanical forces generated during EMT. Nevertheless, it also suggests that the extensive chromatin remodeling associated with EMT is largely insensitive to this mechanical perturbation, highlighting the robustness of the EMT regulatory program. Thus, although mechanical forces can modulate chromatin accessibility, their effects appear to be restricted to a specific subset of genomic regions rather than broadly altering the EMT-associated chromatin landscape.

Collectively, our data indicate that the nucleus responds to compression through a spatially patterned mechanical response: the nucleus forms a robust mechanoprotective shield in the apical region possibly through chromatin closure, while the medio-basal region remains comparatively more permissive to the mechanical regulation of gene expression (Fig. 4H).

## Discussion

Understanding how nuclei control the balance between mechanotransduction and mechanoprotection, particularly during important cellular transitions such as EMT is a fundamental question. Integrating imaging, micromanipulation of specific region of the nuclear envelope while tracking transcription dynamics in living embryos and genomics approaches, we reveal two important features of nuclear EMT mechanics:

1. nuclear mechanical response is spatially patterned within the nucleus.
2. mechanosensitivity is determined by the local chromatin environment rather than by gene sequence, defining a mechanoprotected apical nuclear region and a medio-basal region permissive to force-induced transcriptional responses.

Our results challenge the concept of “mechanosensitive genes” and instead support the idea of mechanosensitive nuclear microenvironments, revealing an apico-basal polarization of the mechanical response.

In the most apical nuclear region, the absence of transcriptional response indicates that the apical nuclear environment serves as a protective role. Given the significant mechanical challenges cells face during EMT (such as epithelial exit and subsequent migration), the establishment of such a mechanoprotective shield may be critical for maintaining a robust transcriptional program, for example preventing the precocious activation of mesodermal differentiation genes. This protection could be mediated by the enrichment of heterochromatin in the apical nuclear region. Indeed, heterochromatinization has previously been proposed as a mechanism to buffer mechanical stress in cultured cells subjected to stretch ^7^. In the context of polarized EMT nuclei during mesoderm invagination, we propose that a heterochromatin-mediated mechanical shield may extend into the underlying euchromatic compartment, thereby rendering apically positioned genes, such as *snail*, less responsive to incoming mechanical stimuli (Fig. 4H).

Our data are consistent with recent work in *Drosophila* testes showing that satellite DNA clustering into chromocentres generates condensate-like assemblies that enhance nuclear mechanical stability ^33^. Similar structures may therefore contribute to the genome-wide apical mechanoprotection observed in EMT nuclei during invagination and in other contexts beyond EMT. Another mechanism that could be involved in the regionalized response to forces is a change in nuclear crowding, a phenomenon previously observed under osmotic stress or compressive forces. Notably, a reduction of nuclear volume can trigger a rapid response through condensates fusion by liquid-liquid phase separation ^34,35^. Given that nuclear volume remains constant during EMT, we could hypothesize that nuclear deformations could unevenly impact different nuclear regions, resulting in heterogeneous crowding along the apico-basal axis.

Of note however, the apico-basal regionalization identified here does not appear to provide a strict predictive rule for the mechanosensitivity of mesodermal genes. Rather, it suggests a spatial bias, with genes located in medial and basal nuclear regions potentially exhibiting greater responsiveness to mechanical forces. For instance, despite its localization outside the apical region associated with reduced mechanosensitivity, *snail* does not respond to compression at its endogenous position. These observations suggest that transcriptional mechanosensitivity depends on distinct subnuclear microenvironments with varying sensitivities to mechanical forces, which appear to be preferentially distributed along the apico-basal axis rather than organized into strictly defined compartments.

Importantly, because the Rabl configuration (marked by centromere and telomere clustering) is widespread during early embryogenesis and persists at later stages in some species ^26,27,36^, position-dependent mechanosensitivity may represent an evolutionarily conserved principle. This spatial organization could also extend to nuclei arranged into chromosome territories, in which some domains are mechanically shielded while others remain responsive, or to alternative architectures such as the “inside-out” nuclei of the mouse retina, where central heterochromatin is surrounded by peripheral euchromatin ^37^. Such organizations may similarly support spatially regionalized transcriptional responses to mechanical cues.

## Acknowledgements

We are grateful to Claudio Collinet, Romain Levayer, Jean-Léon Maître, Marie-Hélène Verlhac, Cyril Esnault, Corinne Benassayag, Sylvain Derossi, Orestis Faklaris and Pierre Bensidoun for their constructive comments on the manuscript. We thank Hadi Boukhatmi for sharing *zfh1* smFISH probe. We acknowledge the Bioinformatics platform BigA, the Imaging platform LITC and the *Drosophila* facility from the CBI, the Montpellier Ressources Imagerie facility (France-BioImaging), the MGX sequencing platform in Montpellier and the Biocampus *Drosophila* facility of Montpellier. MS’s research is supported by the Center National de la Recherche Scientifique (CNRS), the University of Toulouse and funded by grants from the National Agency of Research (ANR, PRC AAPG2021, CellPhy), the Research Association against Cancer (ARC, Programme Labélisé AAP2020, ARCPGA12020010001154_1591) and the Medical Research Fundation (FRM, Equipes FRM, EQU202403018134). Research in the laboratory of M.L. is supported by the Center National de la Recherche Scientifique (CNRS), the University of Montpellier and is funded by grants from the Fondation Bettencourt Schueller, the European Molecular Biology Organization Young Investigator program (EMBO YIP), ERC (LightRNA2Prot), and ANR CellPhy. M.L, A.Z, M.S, T.M1, C.R and A.T. are sponsored by the CNRS; R.B and P.G.I by the FRM.

## Author’s contributions

**Experimental Design, Data Analysis, Method Development and Execution**: ***Ronan BOUZIGNAC***: Led nuclear envelope imaging and characterization; performed optical tweezers (OT) experiments (with Tatiana MERLE and Thomas MANGEAT); conducted compression experiments and quantifications; analyzed genome wide data (with Amal ZINE EL AABIDINE and Lauriane FRITSCH). ***Amandine PALANDRI***: Designed and executed genome wide data. ***Lauriane FRITSCH***: Designed, set up, and executed genome wide experiments. ***Tatiana MERLE***: Set up and performed optical tweezers experiments. ***Amal ZINE EL AABIDINE***: Performed bioinformatic analysis of genome wide data. ***Thomas MANGEAT***: Set up and provided training for optical tweezers experiments and analysis. ***Mélanie GRACIA***: Validated genome wide data; established *sna*- and *twi*-MS2 baselines; performed H2B super-resolution imaging. ***Cyril ANDRIEU***: contribute to quantifications and imaging. ***Pablo GARCIA-IDIEDER***: Generated *sna*-MS2 and *twi*-MS2 lines at different nuclear locations. ***Antonio TRULLO***: Developed automated methods for quantifying MS2 dots at transcription sites (TS) in complex tissues. ***Martine CAZALES***: Set up compression experiments and provided live imaging training. ***Simon MARQUES-PRIETO***: Validated genome wide data. ***Virginia PIMMETT***: Generated and validated the *sna*- and *twi*-MS2-KI line; assisted in probe design and live imaging training. ***Mélissa COSTES***: Assisted Mélanie GRACIA in confirming genome wide data. ***Christian ROUVIERE***: Developed automated methods for analyzing nuclear shape and TS subnuclear localization. **Writing-Editing**: ***Ronan BOUZIGNAC, Mounia LAGHA & Magali SUZANNE***. **Conceptualization, Supervision, and Funding: *Mounia LAGHA & Magali SUZANNE*.**

## Methods

### Experimental animals

The animal model used in this study is *Drosophila melanogaster*, in the context of *in vivo* and *ex vivo* experiments. To adhere to ethical principles, adult flies were anesthetized using CO₂ prior to any manipulation. To prevent accidental release outside the laboratory, deceased flies were frozen before disposal. Stocks of live flies were maintained in incubators at either 18 °C or 25 °C to ensure optimal conditions. The fly food composition included water, agar (0.8%), sugar (4%), flour (7.4%), yeast (2.8%), Moldex (1%), and propionic acid (0.3%). Genotypes and developmental stages used are detailed below. Experiments were performed on both male and female flies without distinction.

### Drosophila Stocks

*The* w*[\**] strain was used as the wild-type control. His2Av-mRFP (Histone H2A variant fused to RFP, from BDSC_23651) were obtained from the Bloomington Drosophila Stock Center (BDSC). To monitor transcription in living embryos, we used the w*[\**]; sqh*-*RFPtKI[3B]; *nos*>MCP-eGFP. The following MS2 reporters were used (always paternally provided): *snail*-MS2 knock-in ^22^, *twist*-MS2 knock-in ^23^, the *snail*-yellow-MS2 BAC ^24^; twi-MS2 transgenes ^28^, sna-MS2 transgenes (generated in this study). For live imaging of the nuclear envelope, we used Nup107-GFP (BDSC_35514) and a Lamin Dm0–TagRFPt knock-in line ^40^. All crosses and embryo collections were carried out at 25 °C.

### sna-MS2 transgenes

The *sna*-MS2 reporter was generated using the minimal regulatory region of the distal enhancer, as described in ^41^, and supplemented with a ParS2 array. This construct will be described in detail elsewhere (Idieder et al., in preparation).

### Twi-MS2 transgenes

The *twi*-MS2 reporter was already described ^28^. In short, it includes the twist upstream cis-regulatory region comprising the distal and proximal enhancers to 50 bp downstream of the transcriptional start site.

### Immunohistochemistry

Embryos were collected after a 2-hour egg-laying period, followed by an additional 2-hour incubation. They were then dechorionated in a 50% bleach solution for 2 minutes. Fixation was performed for 5 minutes in a heptane:formaldehyde (37%) solution (1:1) at 4 °C using a rotary agitator. Embryos were washed twice in PBS containing 0.3% Triton X-100 (PBT). Selected embryos were then sliced using a microscalpel (Micro Knives, Plastic Handle 10315-12, Fine Science Tools).

Embryo slices were blocked in PBT supplemented with 0.5% bovine serum albumin (BBT), followed by overnight incubation at 4 °C with primary antibodies diluted in BBT: mouse anti-Lamin (1:10, ADL67.10, DSHB). After washes in BBT, slices were incubated for 4 hours at room temperature with goat anti-mouse Alexa Fluor 488 (1:100, Interchim). When necessary, Rhodamine-Phalloidin (1:200, Fisher Scientific) was also included.

Samples were then washed in PBT and mounted between two coverslips using a 120 µm deep spacer (Secure-Seal™, Sigma-Aldrich) in Vectashield mounting medium containing DAPI (Vector Laboratories).

### smiFISH

Custom smiFISH probes were designed to target the coding sequences (CDS) of genes of interest using the Stellaris® RNA FISH Probe Designer (Biosearch Technologies, Inc., Petaluma, CA), available online at https://www.biosearchtech.com/stellarisdesigner. Each probe was coupled with a FLAP sequence (X or Y, depending on the probe), as described in ^42^. The smiFISH-FLAP probes were synthesized by Integrated DNA Technologies (IDT).

Embryos were collected, fixed, and sliced as described above for immunohistochemistry. Embryo slices were first washed with PBS containing 1% Tween-20, followed by a 1:1 mixture of PBS-Tween-20 and Wash Buffer A (20% Stellaris RNA FISH Wash Buffer A [LGC Biosearch Technologies, Cat# SMF-WA1-60], 70% H₂O, and 10% deionized formamide).

Samples were then incubated for 1 hour at 37 °C in the dark in Hybridization Buffer (10% deionized formamide and 90% Stellaris RNA FISH Hybridization Buffer [LGC Biosearch Technologies, Cat# SMF-HB1-10]). Embryo slices were incubated overnight in the same Hybridization Buffer containing annealed smiFISH probes (50 nM per oligo).

smiFISH probes were prepared by duplexing 40 pmol of target-specific probes with 100 pmol of FLAPX-Cy3 or FLAPY-Cy5 oligonucleotides in annealing buffer (1× NEBuffer™ 3.1). Duplexing was carried out with the following temperature program: 3 minutes at 85 °C, 3 minutes at 65 °C, and 5 minutes at 25 °C. Probes were then kept on ice until use.

Following overnight hybridization, embryo slices were washed at 37 °C first with Hybridization Buffer, then with Wash Buffer A, and finally with PBS-Tween at room temperature. Immunohistochemistry was performed afterward if required.

### Image Acquisition of fixed samples

Fixed samples were imaged using an LSM900 inverted laser scanning confocal microscope (Zeiss), using a 40x/1.3 oil objective and a piezo stage. Z-stacks were acquired using the laser scanning in High-Resolution mode (Airyscan). Airyscan Z-stacks were processed in ZEN software using the automatic strength (default) and the 3D method.

### Compression experiment

#### Manual compression

For compression experiments, embryos were collected after a 2-hour egg-laying period, followed by an additional 2-hour incubation. The embryos were then dechorionated by immersion in a 50% bleach solution for 2 minutes.

Following dechorionation, embryos were transferred to an agar plate (20 g/L) and covered with Halocarbon Oil to prevent dehydration and facilitate staging; they could be maintained in these conditions for up to 2 hours.

To ensure stability during compression, cover slips were fixed to a metallic frame to prevent bending or any form of movement of the cover slips. Glue prepared by incubating Scotch™ double-sided tape in heptane overnight and placed in the center of the cover slip. One Embryo was gently rolled on the agar surface (avoiding oil) to allow a proper adhesion, and then glued to the prepared cover slip. A drop of 1 µL of Halocarbon Oil was then added to cover the embryo. Samples were imaged using a Zeiss LSM880 confocal microscope in Fast Airyscan mode with Plan-Apochromat 40x/NA 1.3 Oil DIC UV-IR M27 objective at 2x zoom. Time-lapse imaging was performed with a frame interval of 15 seconds. A 488 nm laser at 8% laser power with 850 V gain to detect the MCP-eGFP signal and a 561 nm laser at 8% laser power with 850 V gain to detect the Sqh-RFPtag signal. Images were captured as 16-bit, with 568×568 pixel resolution, with each pixel being 0.18 μm in length and width with a z step of 0.35 µm. Airyscan Z-stacks were processed in ZEN software using the automatic strength (default) and the 3D method.

During imaging, the orthogonal view in ZEN software was used to observe the cellularization process through the Sqh signal. When cellularization reached the basal side of the nuclei, a 22×22 mm coverslip was gently dropped on the sample between two time-frames, resulting in compression of one third of the embryo (Fig. S2G). For each experiment the level of compression was quantified by measuring the nuclear aspect ratio before and after compression.

#### Gene positioning

The relative position of genes along the apico-basal nuclear axis was inferred from their distance to the centromere, based on the established Rabl organization of chromosomes in early *Drosophila* embryos. This relationship was validated by comparing mean nuclear positions measured by smiFISH for *zfh1*, *twi*, *T48*, *sna_bac* (*CG8552*), *brd*, *sim*, *sna* and *Ncad* with their corresponding centromeric distances. Consistent with previous observations ^26^, analysis of genes located on chromosomes II (n = 5) and III (n = 4) revealed a linear correlation with a slope of 0.6907 and a y-intercept of 0.1585. The distance to the centromere was calculated differently depending on the chromosomal arm orientation. For left arms (2L and 3L), where the centromere is located at the end of the sequence, the formula used was: (1 – (gene position / chromosome size)) * 0.6907 + 0.1585.

For right arms (2R and 3R), where the centromere is located at the beginning of the sequence, the formula used was: (gene position / chromosome size) * 0.6907 + 0.1585.

Gene position was calculated as the mean coordinate between the start and end of the coding sequence (CDS). The chromosome sizes used in the calculations were: 2L: 23,493,794 bp; 2R: 25,286,936 bp; 3L: 28,090,333 bp; 3R: 32,079,331 bp (data from FlyBase).

In this study, such calculations were performed automatically using a custom Python script. The obtained gene positions were then classified into 10 bins ranging from 1 (apical) to 0 (basal): 1-0.9; 0.9-0.8; 0.8-0.7; 0.7-0.6; 0.6-0.5; 0.5-0.4; 0.4-0.3; 0.3-0.2; 0.2-0.1; 0.1-0. Due to the unequal distribution of genes across these bins, the number of potentially mechanosensitive genes identified in each bin was normalized by the total number of genes present in the corresponding region. The total number of genes per bin was as follows: 1-0.9 = 0; 0.9-0.8 = 59; 0.8-0.7 = 781; 0.7-0.6 = 1528; 0.6-0.5 = 1608; 0.5-0.4 = 1553; 0.4-0.3 = 1773; 0.3-0.2 = 1516; 0.2-0.1 = 761; 0.1-0 = 0.

The normalized values were then converted into percentages, representing the proportion of responsive genes in each nuclear region. This normalization allowed direct comparison between different gene subsets despite differences in total gene number.

To evaluate whether the genomic distribution of selected genes along the apico-basal axis differed from random expectations, a randomization analysis was performed using 100 gene sets of 70 genes randomly selected from the 799 genes belonging to clusters C1 to C4.

The resulting distributions were summarized by calculating the mean distribution across the 100 randomized gene sets.

### ATAC-seq

#### Embryo collection

All embryo collections were performed at 25 °C. Adult females were allowed to lay eggs for 2 h on agar plates supplemented with fresh yeast, followed by a 1-h incubation. Embryos were then collected and dechorionated for 2 min in 50% bleach. Following dechorionation, embryos were transferred to 20 g l⁻¹ agar plates and covered with halocarbon oil to prevent dehydration and facilitate staging. Embryos were maintained under these conditions for up to 1 h. Cellularized or gastrulating embryos were manually selected under a stereomicroscope. To match the timing of the bulk-compression condition (see ‘Bulk compression’), selected cellularized embryos were aligned and maintained on the agar plate for an additional 5 min. Fifty embryos were collected per replicate. Embryos were subsequently transferred to PBT, washed sequentially in PBT and water, snap-frozen in liquid nitrogen and stored at −80 °C until further processing.

#### Bulk compression

For bulk compression, 50 embryos were aligned and oriented on an agar plate (20 g/L) dorsoventrally to undergo compression along the D-V axis.

A 100 µm spacer was made using two layers of Scotch™ Crystal Clear Tape 600 (50 µm thick) on a 24 × 32 mm coverslip. The central part of the two tape layers was removed using a scalpel, leaving two strips of tape approximately 2–3 mm wide along each side of the coverslip. Small openings (1-2 mm) were also cut in the tape on both sides to allow air to enter the spacer, preventing potential hypoxia of the embryos.

After aligning the 25 embryos, they were picked up using another 24 × 32 mm coverslip on which glue had been deposited, maintaining their alignment and orientation. The embryos were then placed in a homemade compression device consisting of two Plexiglas plates with four holes at each corner and screws and nuts to apply pressure. The coverslip containing the 100 µm spacer was placed on top of the coverslip carrying the aligned embryos. The assembly was then placed in the compression device, and pressure was initially applied manually to ensure a final embryo thickness of 100 µm. The compression was then maintained for 5 minutes by tightening the screws.

After 5 minutes of compression, embryos were transferred into Eppendorf tubes containing PBT and were washed successively in PBT and water before being frozen in liquid nitrogen. After freezing, embryos were stored at -80°C.

#### ATAC-seq and library preparation

ATAC-seq was performed using the Diagenode ATAC-seq Kit (Cat. No. C01080002). For each biological replicate, 25 *Drosophila melanogaster* embryos and 10 *Drosophila yakuba* embryos at the cellularization stage were used,, with *D. yakuba* included as a spike-in control.

Cells were dissociated in a dounce homogenizer using NEB buffer (20 mM HEPES, 10 mM KCl, 0.1% Triton X-100, 20% glycerol). Following dissociation, cells were pelleted by centrifugation at 500g for 5 minutes at 4°C and resuspended in 50 µL of Buffer 1 (provided in the kit).

Tagmented DNA was amplified by PCR (10 cycles), and the resulting libraries were purified. Sequencing was performed by Novogene (https://en.novogene.com/) using paired-end 150 bp reads, generating approximately 6 Gb of data per sample. All conditions were analyzed in three independent biological replicates.

### ATAC-seq analysis

ATAC-seq data were pre-processed and analyzed as described in Galouzis et al. ^48^, with minor adaptations, using *Drosophila yakuba* chromatin as a spike-in control for normalization. Spike-in normalization factors were calculated based on the number of reads mapped to the exogenous *Drosophila yakuba* genome. Spike-in–normalized signal tracks (BigWig files) were generated for *D. melanogaster* using bamCoverage with the --scaleFactor option, applying the normalization factor calculated for each corresponding sample. Significantly differential peaks were defined as FDR < 0.05 and a |log2 Fold change| > 1 in DESeq2 ^49^.

### Hierarchical clustering analysis

Differentially regulated loci (accessible regions for ATAC-seq) were considered significant when they displayed an absolute log2 fold change greater than 1 and a p-value lower than 0.05.

Significant loci identified across the different comparisons were combined into a single non-redundant set of differentially regulated loci. For each locus, the corresponding log2 fold change value from each comparison was extracted and assembled into a matrix.

To focus the analysis on the most variable loci, variability was calculated based on the variance of absolute log2 fold change values across the different comparisons. The 800 loci displaying the highest variability were selected for hierarchical clustering.

Hierarchical clustering was performed using the SciPy hierarchical clustering module with Euclidean distance and the average linkage method.

Heatmaps were generated using the Seaborn clustermap function.

### Data Availability

All genomic data used in this study are publicly available at the NCBI Gene Expression Omnibus (GEO) under accession number GSE343907 (https://www.ncbi.nlm.nih.gov/geo/query/acc.cgi?acc=GSE343907).

Access to the dataset requires the token: krgvkukkxbwdfkb.

### Optical Tweezer

#### Set up

A home-built setup was used for optical trapping measurements. The system combines a fluorescence microscope with an active optical trapping system. This trapping system has two components: one to precisely control the position of the optical trap on the sample, and a back focal plane interferometric (BFPi) system to track the position of the trapped object relative to the center of the laser ^50,51^. Furthermore, the use of back focal interferometry in tissue is also unaffected by light scattering due to its position in the Fourier plane of the detector ^52^.

For calibration and the measurements described below, a steering mirror conjugated to the pupil plane of the microscope (Thorlabs FSM75-P01) was used to achieve nanometric laser deflection. A 3-axis piezoelectric stage (Piezo Concept BIO3) was also used to precisely focus on the trapped object.

The optical trapping system was implemented in a conventional inverted fluorescence microscope (Leica DMI6000 B). The objective lens used is a 100x magnification lens with a numerical aperture of 1.4 (Leica HCX PL APO). To optimize the fill factor of the infrared laser to 85% of the objective lens pupil plane, an X4 afocal telescope consisting of two relay lenses is used (Thorlabs ACA254-050-1064 and ACA254-200-1064). The focused diffraction spot at the objective exit had a width of 600 nm at mid-height, allowing for an efficient optical trap.

The microscope is optimized for simultaneous GFP and RFP imaging combined with the infrared laser trap, thanks to a 3-band dichroic mirror (Semrock Di03-R405/488/561/635-t1-25x36). Imaging and optical tweezers were controlled by combining a National Instruments acquisition card with the Inscoper synchronization box. The Inscoper software managed all synchronization steps.

#### Optical Tweezers calibration and rigidity measurements in complex medium

The complete method and robustness of the optical tweezers in living tissue is discussed in detail in a dedicated paper^53^. The method can be summarized as follow: Internal lipid droplets (refractive index ∼1.42 to 1.5, diameter ∼300-600 nm) were used as probes. Optical trapping was performed using a 1064 nm laser at 152mW for wild-type embryos. These trapping parameters did not perturb tissue-scale behavior, as confirmed by normal gastrulation progression following micromanipulation. During experiments, an increase in temperature of between 1.2°C and 1.6°C was estimated. ^54^.

A back focal interferometry is used to measure the relative displacement of the droplet with respect to the center of the optical trap with a precision of 10–15 nm ^53^. Here, the relative displacement is defined as:

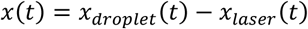

where x_droplet(t) and x_laser(t) are the droplet and laser (trap) positions, respectively.

We calibrated the optical tweezers for each measurement using the fluctuation-dissipation theorem (FDT), with trajectories sampled at 1 kHz and tracked via back focal plane interferometry. This calibration enables quantitative force spectroscopy in heterogeneous environments ^55,56^. We combined passive and active microrheology approaches, applying a 360 nm square-wave displacement at 2.3 Hz, inspired by multiplexed FDT methods ^57^. A 7-second trajectory recording allowed simultaneous estimation of the trap stiffness (kt), the detector conversion factor (β, V/nm), and the effective stiffness of the surrounding medium (k_m). The stiffness of the optical trap was estimated with an absolute error of 10% using the following expression from the fluctuation dissipation theorem, which accounts for all harmonics from the square wave excitation.

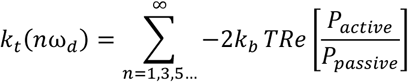

where *P_active_* is the active power spectrum and *P_passive_* is the passive power spectrum. The high-frequency analysis and the number of harmonics provide a very robust estimation of *k_t_*. The mechanical response of the medium can be reconstructed independently at each harmonic frequency once *k_t_* is known. A measurement/calibration step is performed at 1.5 µm from the nucleus, then the procedure is repeated in contact with the nucleus using the same droplet. Contact with the nucleus is checked using both the response curve measured with the back focal plane (BFP) and the small deformation of the nucleus observed in fluorescence images. If the force applied to the droplet is

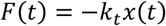

The mechanical response function of the nucleus relative to the BFP is :

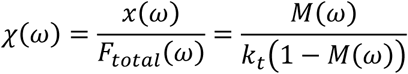

where x(ϖ) is the Fourier transform of x(t), F_total_(ϖ) is the Fourier transform of all contributing forces (e.g. tweezers, nuclear envelope, other contribution) and M(ϖ) is the transfert function defined as:

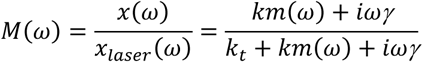

The viscoelastic response of the surrounding medium, km (ϖ) (representing either the nucleus or cytosol, depending on the context) is expressed in pN/μm and is obtained by fitting the experimentally measured transfer function M(ϖ) with a generic polymer-network model:

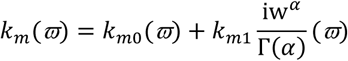

Here, K_mo_ is the dominant elastic component of the nucleus and is the parameter used throughout this work to quantify its stiffness. The second component describes the power-law viscoelastic contribution associated with the actin polymer network. Depending on the experimental conditions, K_mo_ of the nucleus ranges from 10 to 120 pN/μm.

Throughout this work, stiffness is expressed as a complex spring constant (pN/μm) rather than in Pascals. This choice reflects the mesoscopic nature of the measurement, which directly relates the force exerted by the optical trap to the resulting displacement. Note that the link between K_m_(ϖ) expressed in pN/µm and the complex viscoelastic modulus G(ϖ) expressed in Pa is the following equation:

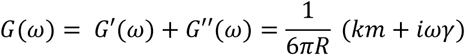

For a droplet radius R of 300 µm and a friction coefficient γ 20 times higher than that of water, our values of G′ for nuclear tension are between 5 and 130 Pa. These values are in agreement to the state of art optical tweezers methods measuring nuclear rheology ^58^.

#### Force induction and transcriptional response

Using the same setup, we deformed nuclei and monitored transcriptional responses with an automated sequence. Nuclei were pre-selected for an active transcription site (TS). The optical trap, initially inactive, was positioned near the nuclear envelope (identified by brightfield contrast; Fig. 4A) using a predefined ROI, with 2×2 pixels binning to enhance signal-to-noise.

The acquisition sequence comprised three phases: (1) **Pre-indentation**: 488 nm LED at 15% power, trap inactive, with 20 Z-stacks (15 slices, 0.4 µm step, 1.5 s/stack); (2) **Indentation**: same LED settings, trap activated with 500 nm square wave modulation at 0.5 Hz, with identical Z-stack acquisition; (3) **Brightfield snapshot**: a single-plane image at the Z-stack center. Successful indentation was confirmed by visual nuclear deformation. We selected experiments where TS intensity was reliably trackable throughout, ensuring the TS remained fully within the Z-stack (xz projection in Fig. 4C-C’).

### Quantification and statistical analysis

#### Nuclear segmentation

Nuclei segmentation was performed using immunofluorescence staining with an anti-Lamin Dm0 antibody. Cellpose ^59^ was configured with the cyto2 model, applying a flow threshold of 0.4 and adjusting the estimated cell diameter per image to optimize results. The resulting labels were then filtered with the MorphoLibJ plugin ^60^ using Label Size Filtering (Greater_Than; Size Limit: 50,000 pixels). The central (middle) plane of each nuclear mask was extracted using a custom macro in Fiji (described in ^52^. Given the variability in nuclear orientation, each nucleus was realigned with the image y-axis to minimize artifacts in aspect ratio and deformation index measurements. Quantification was restricted to nuclei located within the invagination.

#### Nuclear circularity

Nuclear circularity was quantified using the Shape Descriptor option in Fiji. Measurements were performed manually using immunofluorescence staining with an anti-Lamin Dm0 antibody.

Circularity was measured for each nucleus at five different positions along the A-P/B axis of the nucleus. The five positions were defined as 0.9, 0.7, 0.5, 0.3, and 0.1, where 1 corresponds to the apical part of the nucleus and 0 to the basal part.

#### Nuclear aspect ratio and volume

Nuclear aspect ratio and volume were quantified using Label Analyzer (2D,3D) from the SCF-MPICBG plugin in Fiji (Version 1.2.0).

#### Mean local curvature

The local curvature was analyzed only in the apical region and not over the entire nucleus. The upper part of each nucleus was defined using coordinates obtained from the “Label Analyzer (2D, 3D)” of the SCF-MPICBG plugin, based on the center of mass and bounding box. Based on a selection of the upper part of the nucleus, angles were calculated using the Array.getVertexAngle function from Fiji. The mean local curvature for each nucleus was defined as the mean of the top 10% maximum angles. These angles were also used to compare their distribution in *wild-type* nuclei.

#### % Deformed Nucleus

Based on the angles calculated for the deformation index, a nucleus was classified as deformed when it contained at least one angle greater than 60°. This threshold was chosen because it showed the most significant contrast between embryos at the cellularization stage, which predominantly contain undeformed nuclei, and embryos during invagination, which mainly exhibit deformed nuclei (data not shown).

3D nuclear reconstruction Based on the segmented nuclear mask, 3D reconstructions were generated using the Surface tool in IMARIS.

#### smiFISH TS analysis

smiFISH analysis was performed using previously published custom Python software ^61^ available online: https://github.com/ant-trullo/smFiSH_software. From this analysis, we extracted normalized intensity and 3D coordinates of all TS. Then, we determined manually in Fiji, for each TS, if it belongs to a deformed or undeformed nucleus, using the same method described above (% of deformed nuclei), which allows us to quantify the “number of active TS” and the “TS intensity per nucleus” (sum of normalized intensities of all TS within a given nucleus).

### Compression and live transcription

Live transcription was analyzed using a feature (detection and tracking MS2/MCP TS in the absence of a nuclear marker) of a custom Python software previously published in ^23^ available online: https://github.com/ant-trullo/OptoTrack. From this analysis, we extracted all TS with their normalized intensities and 3D positions. Only tracks with a duration of at least 10% of the total imaging time were retained for analysis. The “Intensity Ratio” before and after compression was calculated as the ratio of the mean intensity of all tracks during the 5 minutes (corresponding to 20-time frames) before and after compression.

The “Nucleus deformation” was manually assessed in Fiji, based on the shadow projected by the nucleus in the Sqh signal. 10 nuclei were quantified after and before to calculate the “Nucleus deformation” ratio also use as a readout of the level compression applied for each experiment.

### Transcription response to Force induction

Movies recorded with the trap off and on were concatenated. Only movies in which TS remained entirely within the Z-stack throughout the acquisition were included in the analysis. Prior to analysis, bleach correction and bandpass filtering were applied in Fiji. The same Python-based software used for the compression experiments was then employed for quantification.

The distance between the TS and the optical trap was calculated using the 3D coordinates of the TS extracted from the first frame and the trap position, defined by a manually placed ROI during the experiment and register within the movie metadata.

The “TS Intensity Ratio” was defined as the ratio between the mean normalized intensity of the last 10 frames with the trap off and the mean normalized intensity of the last 10 frames with the trap on. For each experiment, a TS from an unmanipulated nucleus was used as a control.

#### Optical Tweezers: Relaxion

Nuclear relaxation was assessed by comparing the nuclear area in 2D before and after the indentation phase. A nucleus was considered to have undergone complete relaxation if its area after indentation (end of the movie) was at least 95% of its area before indentation (beginning of the movie). Values below this threshold were classified as partial relaxation. Note: none of the indentations were maintained. Nuclear areas were manually measured using Fiji.

#### Heterochromatin analysis

3D Heterochromatin analysis was performed using the DAPI staining. Cellpose configured with the cyto2 model was used for segmentation, applying a flow threshold of 0.4 and adjusting the estimated cell diameter per image to optimize results. Volume and total intensity were quantified using Label Analyzer (2D,3D) from the SCF-MPICBG plugin in Fiji (Version 1.2.0). Total intensity values were normalized to the mean DAPI signal of the euchromatin.

#### Lamin analysis

A Lamin-RFP knock-in line was used to visualize the lamin signal along the nuclear envelope at the nuclear mid-plane. A custom Fiji macro was developed to extract the signal around the entire nuclear contour. Apical and basal regions were defined as the top 25% and bottom 25% respectively, relative to the mean vertical position of each nucleus. The apical-to-basal intensity ratio was then calculated and plotted.

---

**Figure S1:**
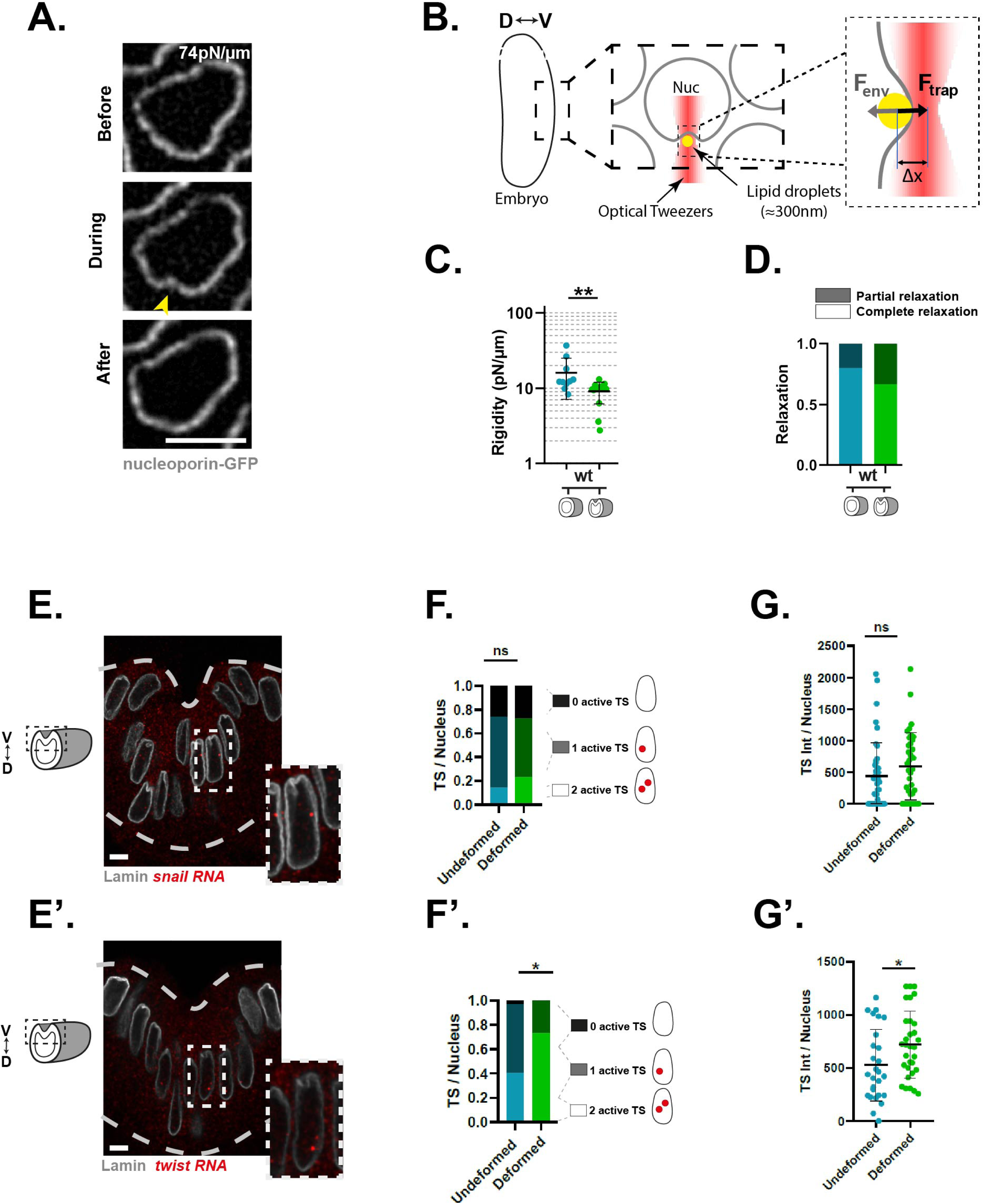
Nuclear stiffness and sna and twi nascent transcription at the onset of EMT. Related to Fig. 1. **A.** Time-lapse snapshots from optical tweezer experiments showing 3 successive time points: before indentation (calibration, top), during indentation (measurement, middle), and after indentation (relaxation, bottom), in *wt* nuclei at 74pN/µm. **B.** Schematic representation of the optical tweezer method using lipid droplets as internal probes in living embryos. F_env_ = restoring force of the nuclear envelope (resulting from its rigidity); F_trap_ = optical trap force; Δx = distance between the lipid droplet and the center of the trap. Knowing F_trap_ and Δx, we can estimate F_env_. **C.** Quantification of nuclear rigidity measured by optical tweezers, 10 minutes before (n1) and at the onset of EMT (n2) ( N=8, n1=10, n2=14). **D.** Quantification of nuclear envelope relaxation 7 sec after the release of optical tweezer-induced deformations, for the same conditions as in C (N=8, n1=10, n2=19). **E.** Transverse sections of embryos stained for Lamin Dm0 and hybridized via smiFISH for *snail* (K) or *twist* (K’). Dashed lines indicate the apical and basal sides of the tissue. **F.** Quantification of the number of TS per nucleus or transcriptional activation for *snail* (L) and *twist* (L’) in undeformed and deformed nuclei of *wt* embryos (for *snail*: undeformed N=8, n=43; deformed N=8, n=37; for *twist*: undeformed N=6, n=26; deformed N=6, n=32). **G.** Quantification of the normalized total TS intensity or transcriptional intensity per nucleus for *snail* (N) and *twist* (N’) in undeformed and deformed nuclei of *wt* embryos (for *snail*: undeformed N=8, n=43; deformed N=8, n=37; for *twist*: undeformed N=6, n=26; deformed N=6, n=32). Statistical significance was assessed by the Mann−Whitney test (Fig. S1 C, G, G’), Chi Square Test (Fig. S1F, F’). (**\***=p<0.05; **\*\***=p<0.01; **\*\*\***=p<0.001;****=p<0.0001). N=embryos, n=nuclei. Scale bar = 5 μm.

**Figure S2:**
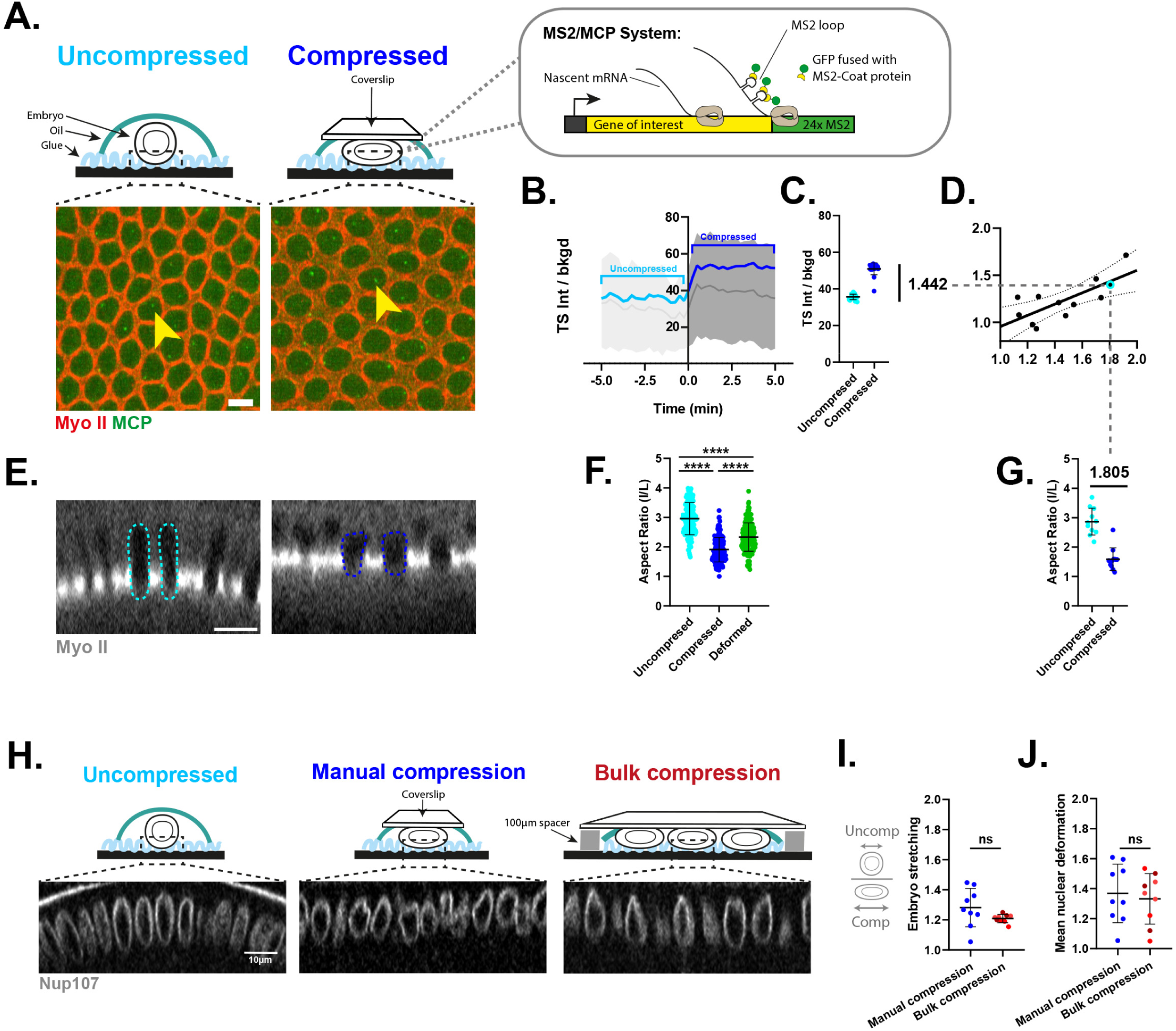
Embryo compression methodology. Related to Fig. 2. **A.** Schematic representation of the mechanical compression setup. For a detailed description, see Methods. Time-lapse snapshots of *twist-MS2*-*KI* embryos before (left) and after (right) compression. Yellow arrows indicate the same TS over time. MyoII is shown in red, MCP-eGFP (TS) in green. **B.** Representative traces of mean normalized TS intensities over time during compression for *twist-MS2-KI* embryos. Light blue indicates before compression, dark blue after compression. The bracket along the x-axis shows the time window used to quantify the TS intensity ratio shown in the scatterplot in C. **C.** Mean TS intensity before and after compression based on the representative traces of normalized TS intensities in panel B. **D.** Curves showing the variation in TS intensity (ratio before vs after compression) in relation to the level of nuclear deformation in *twist-MS2*-*KI* embryos. **E.** Orthogonal views of embryos before (left) and after (right) compression, MyoII is shown in white. Dashed outlines highlight nuclear edges before (light blue) and after (dark blue) compression. **F.** Quantification of nuclear aspect ratio before and after compression, compared with naturally deformed nuclei in control embryos (before/after compression: N=17 n=170, deformed: N=13, n=223). **G.** Quantification of nuclear aspect ratio before and after compression based on nuclei shown in panel E (before: n=10 nuclei; after: n=10 nuclei). Note that we use the variation of the nuclear aspect ratio as a readout of nuclear deformation and the level of compression applied during a given compression experiment. **H-J.** (H) Images showing nuclear aspect in embryos either uncompressed (left), manually compressed (middle) or upon bulk compression (right), (I) with the corresponding quantification of embryo stretching (Manual N=9 n=9, Bulk N=3 n=9), (J) and nuclear aspect ratio (Manual N=9 n=9, Bulk N=3 n=9). Statistical significance was assessed by the Mann−Whitney test, (**\***=p<0.05; **\*\***=p<0.01; **\*\*\***=p<0.001 ****=p<0.0001). N=embryos, n=nuclei. Scale bar = 5 μm

**Figure S3:**
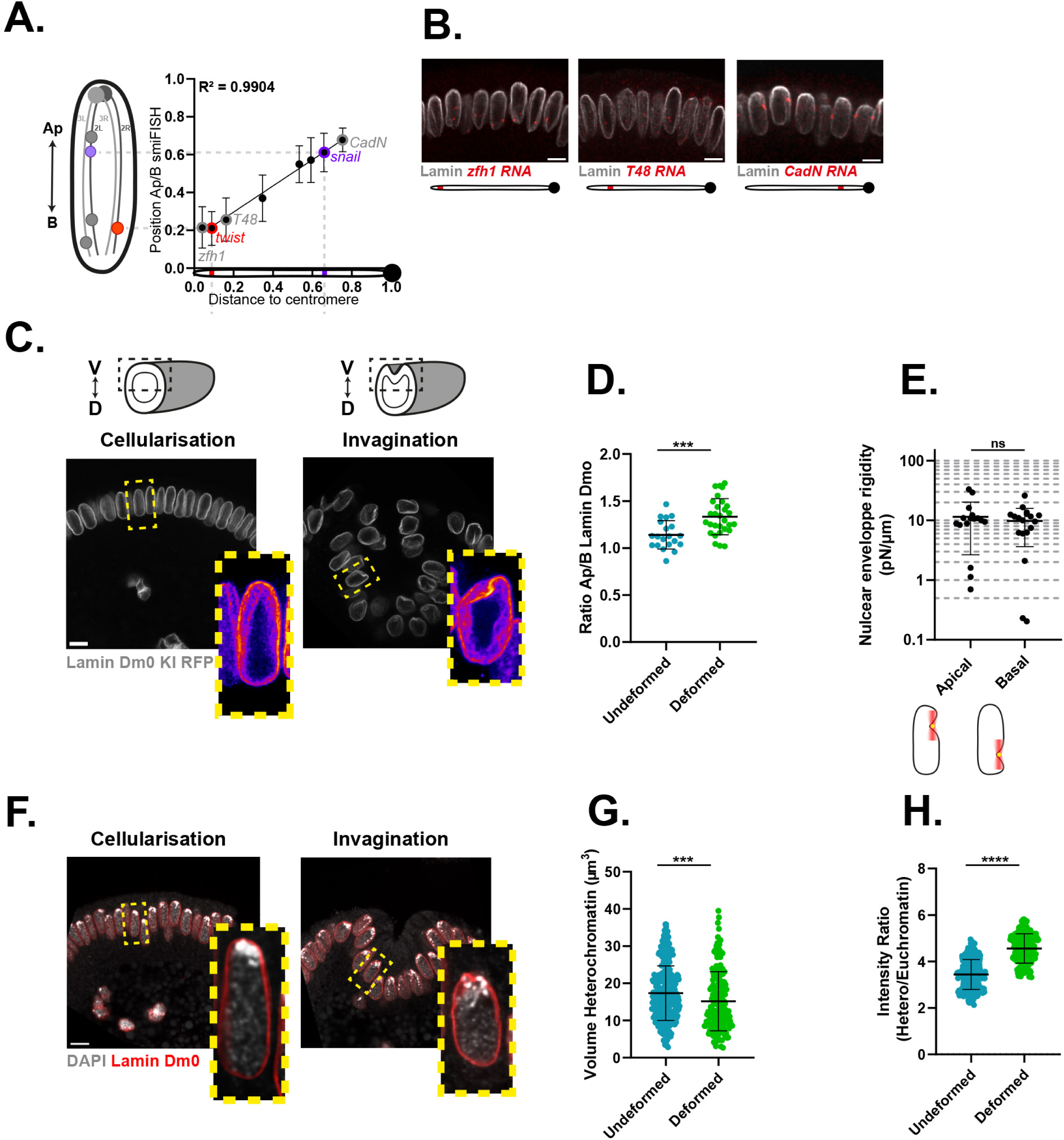
Apico-basal heterogeneity in nuclear mechanics. Related to Fig. 3. **A.** Correlation between gene positioning along the apico-basal axis of the nucleus and their genomic distance to the centromere. *snail* (violet), *twist* (red) and *zfh1*, *T48*, *cadN* (grey). **B.** Transverse sections of embryos stained for Lamin Dm0 and hybridized via smiFISH for *zfh1* (left), *T48* (middle) and *cadN* (right). **C.** Cross-sections of *LaminKIRFP* embryos during cellularization (pre-EMT, left) and invagination (EMT, right). Insets show selected nuclei using the Fire LUT to visualize signal distribution. **D.** Quantification of the apical-to-basal Lamin Dm0 intensity ratio in undeformed and deformed nuclei (undeformed: N = 2, n = 20; deformed: N = 3, n = 29). **E.** Quantification of nuclear rigidity at two positions along the lateral nuclear envelope, as indicated in the schematics, using optical tweezers in *wt* embryos. Measurements were performed at apical and more basal positions (N = 6; n=16), with both measurements acquired 5 µm apart within each nucleus. **F.** Cross-sections of *w* embryos stained for Lamin Dm0 and DAPI during cellularization (pre-EMT, left) and invagination (EMT, right). **G.** Quantification of heterochromatin volume based on DAPI-dense regions in undeformed and deformed nuclei (undeformed: N = 3, n = 284; deformed: N = 3, n = 172). **H.** Quantification of total heterochromatin intensity, normalized to mean nuclear DAPI intensity, in undeformed and deformed nuclei (undeformed: N = 3, n = 282; deformed: N = 3, n = 172). Statistical significance was assessed by the Mann−Whitney test (**\***=p<0.05; **\*\***=p<0.01; **\*\*\***=p<0.001). N=embryos, n=nuclei. Scale bar = 5 μm

## Notes

### Competing Interest Statement

The authors have declared no competing interest.

### Summary of Updates

The present study includes substantial new data (in particular data from figure 3 and 4) that change its central conclusion. In particular, the demonstration that nuclear position governs mechanosensitivity, which is the principal finding of the current manuscript, was absent from the earlier submission.

